# Disassortative mating in replicated secondary contact experiments in nature

**DOI:** 10.64898/2026.04.28.721371

**Authors:** Lucas Eckert, Daniel I. Bolnick, Catherine L. Peichel, Andrew P. Hendry, Rowan D.H. Barrett

**Affiliations:** Department of Biology, McGill University, Montréal, QC, Canada; Department of Ecology and Evolutionary Biology, University of Connecticut, Storrs, CT, USA; Division of Evolutionary Ecology, Institute of Ecology and Evolution, University of Bern, Bern, Switzerland

**Keywords:** ecological speciation, allopatric speciation, reproductive isolation, assortative mating, disassortative mating, mate choice, secondary contact, threespine stickleback

## Abstract

Ecological speciation is now regarded as one of the primary processes by which new species are generated. Adaptive divergence in allopatry begins this process, but it is often unclear when and how mechanisms that promote reproductive isolation, such as assortative mating and selection against hybrids, evolve. Here, we test for evidence of these mechanisms across replicated secondary contact experiments in natural settings. We introduced four to eight allopatric populations of threespine stickleback (*Gasterosteus aculeatus*), in both single-ecotype and mixed-ecotype treatments, into nine natural lakes, after which we inferred mating patterns by genotyping the resulting F1 generation. Contrary to expectations from the literature, we found no evidence of assortative mating or partial reproductive isolation among the introduced source populations. Instead, we detected evidence of disassortative mating by source population in three lakes and some evidence of disassortative mating by ecotype in one lake. These mating patterns were both context-dependent and population-dependent, varying substantially across lakes receiving the same source populations, and with some source populations generally displaying greater tendencies for disassortment. The absence of positive assortative mating in any replicate demonstrates that adaptive divergence in allopatry alone might be insufficient to generate assortative mating in many cases, while the possibility of disassortative mating in these contexts poses an additional hurdle on the path toward speciation.

## Introduction

Ecological speciation is now regarded as one of the primary processes by which new species are generated (Nosil, 2012; Rundle & Nosil, 2005; Schluter, 2009). This process often begins in allopatry, where divergent environmental conditions impose divergent selection, leading populations to evolve locally adaptive phenotypes. Over time, such adaptive divergence may give rise to mechanisms that reduce gene flow and promote reproductive isolation among populations (Rundle et al., 2000). Although the initial step of this process, adaptive divergence, is well established and reflected in the widespread occurrence of local adaptation (Hereford, 2009; Leimu & Fischer, 2008), it remains unclear under what conditions adaptive divergence translates into reproductive isolation, and when the mechanisms that facilitate this transition begin to evolve (Sobel et al., 2010).

Ecological speciation often begins with a period of allopatry, but progress toward reproductive isolation only becomes evident following secondary contact. Secondary contact however, not only reveals such progress but might also be critical in reinforcing the mechanisms promoting reproductive isolation (Kirkpatrick, 2001; Rundle & Schluter, 1998). During secondary contact, hybrid offspring can express intermediate or transgressive phenotypes that perform poorly in the current environment, resulting in selection against hybrids (Hatfield & Schluter, 1999; Hendry et al., 2009; Thompson et al., 2024). Such selection can maintain phenotypic and genetic divergence between populations (DiVittorio et al., 2020; Gow et al., 2007) and provides a context in which assortative mating (preferential mating with individuals from the same population or with similar phenotypes) can be favoured. By reducing the production of low-fitness hybrids, assortative mating maintains local adaptation and incidentally contributes to reproductive isolation (Hollander et al., 2005; Liou & Price, 1994). Because secondary contact both exposes and contributes to speciation, it is often unclear how much reproductive isolation arose prior to secondary contact, from divergence in allopatry alone. Direct tests of this question are rare, as the onset and early stages of secondary contact among previously allopatric populations are seldom observed in nature.

Threespine stickleback (*Gasterosteus aculeatus*, stickleback hereafter) have proved a powerful system for studying ecological speciation, owing to wide variation in the strength of reproductive isolation observed among populations and ecotypes (Hendry et al., 2009; Lackey & Boughman, 2017; McKinnon & Rundle, 2002). For example, in the sympatric benthic-limnetic species pairs (Schluter & McPhail, 1992), hybrids experience reduced fitness (Hatfield & Schluter, 1999; Thompson et al., 2022), and positive assortative mating based on morphology is well documented (Bay et al., 2017; Conte & Schluter, 2013; Head et al., 2013; Nagel & Schluter, 1998). Assortative mating has also been reported in some parapatric systems, such as divergence between lake and stream populations (Berner et al., 2017; Eizaguirre et al., 2011), although assortment is generally weaker and less consistent than in sympatry (Raeymaekers et al., 2010; Räsänen et al., 2012). In contrast, mating patterns among allopatric populations have received comparatively little attention (Vines & Schluter, 2005), though a recent experimental study in semi-natural ponds provided evidence for assortative mating among ecotypes originating from populations that evolved with and without sculpin (Roesti et al., 2025). Further investigation of mating patterns among allopatric populations are needed to clarify the specific mechanisms and chronology underlying the evolution of reproductive isolation.

Taken together, evidence from sympatric and parapatric stickleback systems has led to the expectation that divergent populations should exhibit assortative mating, and the limited evidence from allopatric systems seems to support this notion. This expectation aligns with a broader pattern across taxa, where some degree of assortative mating appears to be common (Janicke et al., 2019; Jiang et al., 2013; Rios Moura et al., 2021). Although the opposite phenomenon – disassortative mating – has been documented (Hedrick et al., 2016, 2018; Maisonneuve, Chouteau, et al., 2021; Takahashi & Hori, 2008), and theoretical models predict contexts in which it should be favoured (Bolnick & Doebeli, 2003; Maisonneuve, Beneteau, et al., 2021), such cases are typically treated as exceptions; consequently, the potential impact of disassortative mating on ecological speciation remains largely unexplored.

Here, we test this expectation of assortative mating by examining mating patterns following experimental secondary contact among allopatric stickleback populations introduced into natural lakes. Our experimental design enables tests of assortative mating across multiple levels simultaneously (population, region, and ecotype) as well as along continuous gradients of morphological and genetic divergence. By leveraging replicated whole-lake introductions, we assess mating patterns in ecologically realistic settings, providing rare insight into the conditions under which reproductive isolation evolves in nature.

## Methods

### Experimental design

Our study leveraged adaptive divergence among allopatric stickleback populations to test for evidence of partial reproductive isolation. Specifically, we focus on variation along the benthic-limnetic continuum, representing ecological divergence among populations of lake-dwelling stickleback (Bolnick & Ballare, 2020; Matthews et al., 2010; Schluter & McPhail, 1992; Willacker et al., 2010). Divergence along this axis has been linked to lake bathymetry, through its influence on prey community structure and abundance, which in turn influences the trajectory of stickleback adaptation (Schluter & McPhail, 1992; Willacker et al., 2010). Populations of the benthic ecotype are typically found in small shallow lakes, where they feed predominantly on benthic macroinvertebrates, while populations of the limnetic ecotype are typically found in large deep lakes where they feed on limnetic zooplankton. There is well-defined, but continuous, phenotypic divergence along this axis, especially in morphology (Bolnick & Ballare, 2020; Haines et al., 2023; Lavin & McPhail, 1986; Schluter & McPhail, 1992; Willacker et al., 2010).

Using this system, we conducted a replicated introduction experiment in natural lakes. In 2018, nine natural lakes on the Kenai Peninsula of Alaska were treated with rotenone by the Alaska Department of Fish and Game (ADFG), thereby eliminating invasive predatory fish and incidentally all other fish life (Couture et al., 2022). In 2019, we introduced stickleback (native to the region) to these lakes to rehabilitate the ecosystems. Each of these lakes (hereafter referred to as the ‘recipient lakes’) received fish from either four ‘benthic’ source lakes, four ‘limnetic’ source lakes, or a mixture of both ecotypes, always in a relatively equal mixture of fish from each source population (Figure 1, Table S1 lake details, Table S2 for stocking proportions). Source populations were selected based on fish morphology by quantifying morphological variation among populations using a linear discriminant (LD) analysis of eight functional traits related to feeding and swimming (Hendry et al., 2024), and selecting populations towards the extremes of the primary (benthic-limnetic) axis of divergence (Figure 1). Details of the design and implementation of this experiment can be found in Hendry et al. (2024). One of the lakes that received both ecotypes (GL) is exceptional for several reasons, most notably with the introduction occurring three years later than the others and receiving only seven of the eight source populations received by the other replicate (details in Supplementary Methods).

**Figure 1:**
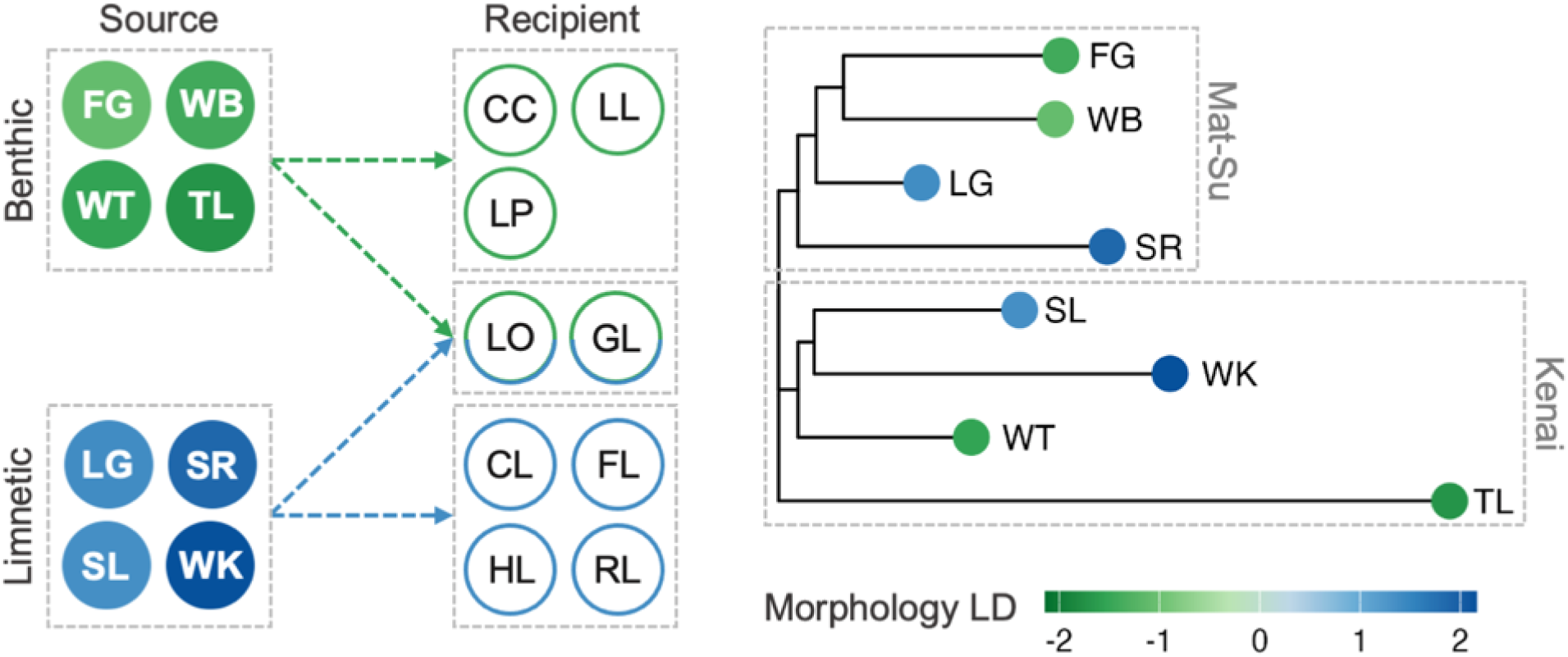
Experimental design: Source lakes are coloured according to their mean morphology linear discriminant (LD) score (Methods for details), and recipient lakes are coloured according to the ecotype of fish they received. The tree on the right was built from neighbour-joining genome-wide F_ST_ between the source populations (Methods for details), again coloured by mean LD score, illustrating the range and combinations of genetic and morphological divergence among populations. The two clusters represent the two geographic regions: the Mat-Su Valley and the Kenai Peninsula.

Several features of this experimental design allow for effectively testing mechanisms contributing to mating patterns. First, two lakes received fish of both ecotypes, allowing us to test if adaptation along the major axis of ecological divergence has resulted in progress toward reproductive isolation. Further, multiple populations of each ecotype were introduced into each recipient lake, allowing us to separate the effects of assortment by ecotype from assortment by population. Second, considerable phenotypic and genetic variation exists among populations within each of these ecotypes (Haines et al., 2023; Hendry et al., 2024) and many traits evolve independently of benthic-limnetic divergence (Bolnick et al., 2026; Neumann et al., 2025), providing opportunity for assortment by population, even in the lakes that received only a single ecotype. Finally, we chose source populations from two geographic regions, the Matanuska-Susitna (Mat-Su) Valley and the Kenai Peninsula, with two populations of each ecotype from each region, and genome-wide F_ST_ among populations primarily reflects isolation by distance (Figure 1). The factorial crossing of ecotype and region separates the main axis of adaptive divergence from overall genetic similarity, allowing us to independently test how each of these factors contribute to mating patterns.

### Sampling and DNA extraction

We sampled 100 fish from each recipient lake both one year and two years post-introduction, in late May of each year, using unbaited Gee’s (G40 – ¼” mesh) minnow traps. All lethal collections were conducted with IACUC approval (University of Connecticut: A19-015, A22-006) and permits from the State of Alaska (SF2020-103d, P-21-012). We euthanized fish on site with buffered tricaine methanesulfonate (MS-222), measured standard length with digital calipers (to 0.01mm), weighed the mass of each fish on a digital scale (to 0.01g), and took a tissue sample from the caudal fin for DNA extraction. We used a standard phenol-chloroform DNA extraction protocol; tissue samples were incubated at 55°C in digestion buffer with proteinase K, before the DNA was isolated using an isoamyl-phenol-chloroform solution and precipitated and washed in ethanol.

### Genotyping and ancestry inference

To explore mating patterns, we inferred the ancestry of fish from the recipient lakes to determine the parental source populations. To infer ancestry, individual fish were genotyped for single nucleotide polymorphisms (SNPs) specific to each source population, identified through pool-seq data originally collected by Weber et al. (2022). After designing and validating the genotyping arrays, the final set of SNPs was 158 in total (20 SNPs per source population on average) with an average allele frequency of 81% in the respective source population. Ancestry proportions were assigned to each fish based on the numbers and frequencies of source-specific SNPs identified in each individual from the genotyping data. Ancestry proportions were rounded to the nearest fraction possible according to the maximum potential generations (assuming the fastest generation time to be one year). For example, in the F1 generation, which we focus on in this study, all ancestry proportions must be either 0.5 (F1 hybrid between two populations) or 1 (F1 offspring of a cross between parents from the same population).

The accuracy of the above ancestry inferences was investigated through simulations in R (v4.5.1), parameterized with the number and frequency of the SNPs in our genotyping arrays. These simulations confirmed very high accuracy in the F1 generation, with an error rate (the proportion of fish for which we fail to identify one of the parental source populations) of <0.001%. Methods for genotyping and ancestry inference are further detailed in Eckert et al. (2026), and have been used in other several other studies of this system (Bolnick et al., 2026; Heckley et al., 2026). Ancestry inferences also have been corroborated by independent data (SNP variant calling from 3’TagSeq transcriptomes; Runghen et al., unpublished data). Using these genotyping arrays, we genotyped 95-97 fish from each of the recipient lakes both one year and two years post-introduction, except for GL (G Lake); given its out-of-sequence introduction, we genotyped 190 fish, but only from two years post-introduction.

### Age class inference

Subsequent analyses include only fish from the F1 generation, such that we can confidently infer mating patterns among the parental source populations. Identifying these fish is made challenging by the presence of overlapping generations, as breeding can occur at one or two years of age. To discriminate between generations, we used information from both genotyping data and measurements of standard length for each fish (see Supplementary Methods). Incorporating both streams of data, we omitted individuals that were likely to be from the parental or F2 generations, leaving us with just a sample of fish from the F1.

### Tests of assortative mating

To quantify the degree of assortative or disassortative mating by source population in each recipient lake, we determined if pure or hybrid genotypes were over- or under-represented compared to expectations of random mating. To make this determination, we treated the overall frequency of each source population in each lake as equivalent to ‘allele’ frequencies, and the frequencies of each pure and hybrid type as equivalent to ‘genotype’ frequencies. Here, the homozygous genotypes represent fish with ‘pure’ ancestry from a single source population, whereas the heterozygote genotypes represent ‘hybrids’ between two source populations. To calculate the expected genotype frequencies, we derived the ‘allele’ frequencies of each source population directly from the F1 offspring rather than the initial stocking frequencies; this is an important distinction as some source populations were initially more successful than others (Eckert et al., 2026). Although we infer mating patterns from these source population frequencies, we acknowledge the possibility that premating isolation might have influenced source population success.

We tested for deviations from random mating by estimating an assortment parameter (*â^s^*) that best fit the observed genotype counts in each lake (Weir & Hill, 2002). This assortment parameter is conceptually analogous to F_IS_, describing the proportional deficit or excess of homozygotes relative to expectations of random mating, though estimated differently than traditional F_IS_ (details in Supplementary Methods), making it more robust towards multi-allelic cases and rare alleles (Weir & Hill, 2002).We estimated this assortment parameter and confidence intervals around it using maximum likelihood estimation and tested for significance by comparison to a null model of random mating (*â_s_* = 0) using a Likelihood Ratio Test (LRT). We supplemented these analyses with traditional population genetic assessments using exact tests of Hardy-Weinberg equilibrium (HWE), and calculating observed heterozygosity (H_o_), expected heterozygosity (H_e_), and inbreeding coefficients (F_IS_) within each lake (Supplementary Methods).

In addition to testing for assortment by source population, we tested for assortment by region (Kenai versus Mat-Su) and by ecotype (benthic versus limnetic). To test for assortment by region, we fit an additional model with both the original assortment parameter (*â_s_*), representing assortment by source population, and an additional parameter (*â_r_*), representing assortment by region (details in Supplementary Methods). Inclusion of both parameters allows us to test if regional assortment significantly explains additional variation when also considering assortment by source population. Again, we estimated both parameters and confidence intervals through maximum likelihood estimation and used a Likelihood Ratio Test to compare to the previous model including just the source population assortment parameter. We used the same framework to test for assortment by ecotype (*â_e_*) in the two lakes that received both ecotypes. Within each type of assortment, we adjusted all p-values from the LRTs to account for multiple testing using a Benjamini-Hochberg correction.

### Source population specific patterns

Individual source populations might contribute differentially to the observed mating patterns due to variation in mating preferences. Further, considerable variation existed in the contribution of each source population to each lake (Eckert et al., 2026), meaning that overall mating patterns could be heavily influenced by strong mating preferences in a single abundant source population. To explore source population specific patterns, we calculated source population specific inbreeding coefficients (F*_i_*) in each lake, analogous to allelic inbreeding coefficients weighted by expected heterozygosity (Nei, 1977), effectively quantifying how each individual source population contributed to the overall estimate of F_IS_ in each lake.

### Morphological and genetic divergence

We next explored if the degree of morphological and genetic divergence among source populations influenced the mating frequency of those populations. Under positive assortative mating, we not only expect an overrepresentation of pure genotypes, but among hybrid genotypes, we might expect overrepresentation of hybrids between parental populations that are morphologically or genetically similar. We tested this expectation using models of the observed count of each hybrid genotype, including the level of morphological or genetic divergence among the parental source populations of that genotype as a predictor. We excluded all pure genotypes from this analysis because their inclusion would yield similar results as the assortment tests as much of the variation in divergence would be explained by the contrast between pure and hybrid genotypes.

To quantify morphological divergence, we extracted the mean linear discriminant (LD) score for each source population from Hendry et al. (2024). These LD scores were based on linear measurements of eight traits, representing variation in body shape and trophic morphology, using a linear discriminant analysis to find the axis of variation that best discriminates among ecotypes. The morphological divergence between two populations was simply the absolute difference in population mean LD scores on this axis (Table S3). To quantify genetic divergence, we estimated two different metrics of pairwise F_ST_ between source populations: one across the entire genome (genome-wide divergence) and one across putatively adaptive windows in the genome (adaptive divergence). Genome-wide genetic divergence was estimated across the autosomes, while adaptive genetic divergence was estimated across regions that we putatively associated with adaptation along the benthic-limnetic divergence (Supplementary Methods). Although the regions we identify only capture one specific axis of adaptation, it should parallel the major axis of morphological and ecological variation among these populations. Details for data processing, filtering, and F_ST_ estimation are in the Supplementary Methods and the resulting F_ST_ estimates are in Table S3.

We used generalized linear models for each recipient lake, modelling the observed genotype counts using a Poisson error distribution with a log link, including the log of the expected genotype count under Hardy-Weinberg equilibrium as an offset. We included the level of either morphological or genetic divergence of the parental source populations of that genotype as a fixed effect, fitting separate models for each type of divergence, assessing significance using Wald tests, and adjusting p-values within each type of divergence with a Benjamini-Hochberg correction. We also tested each type of divergence across all lakes, as a mixed model with lake as a random effect.

### Selection on hybrid individuals

Since we are inferring mating patterns from the offspring after one or two years, differential viability or survival among genotypes could contribute to the observed patterns. We explored this possibility through two approaches, neither of which are conclusive, but either of which could potentially elucidate strong selection. First, we used standard length at a given age and body condition (Fulton’s condition factor) as proxies for fitness (Fletcher & Wootton, 1995; Wootton, 1973; Wootton & Evans, 1976) and compared between pure and hybrid fish in each lake, though we acknowledge that these proxies are imperfect. To test for differences between genotypes, we fit linear models for each lake separately with either standard length or body condition as the response variable and genotype (hybrid or pure) as a fixed effect, along with age for models of standard length. In the two lakes with both ecotypes, we also compared standard length between pure and hybrid genotypes by ecotype; though we did not compare condition among ecotype hybrid classes, as divergence in body depth among ecotypes results in benthic fish generally having more mass per unit length. Second, we tested if the relative proportion of pure versus hybrid fish in the F1 generation changed from year one to year two, which would be indicative of differential survival among genotypes during that time span. Specifically, we used Fisher’s exact tests on the proportions of hybrid fish to test for differences between the first and second year of sampling.

## Results

### Tests of assortative mating

#### Assortment by source population

We found considerable variation in mating patterns across lakes, with evidence of disassortative mating by source population in some lakes and random mating in others (Figure 2; Table S4). Evidence of disassortative mating was found in three lakes: FL ( = −0.13, LR = 9.87, p_adj_ = 0.015), RL ( = −0.11, LR = 7.41, p_adj_ = 0.029), and GL ( = −0.08, LR = 5.76, p_adj_ = 0.049). In a fourth lake (LP), a tendency for disassortative mating was also present ( = −0.05), although this was not significant after correcting for multiple testing (LR = 4.12, p_adj_ = 0.095). In three of these cases (RL, LP, and GL), the maximum likelihood estimation lay at the lower confidence interval boundary (Figure S5). This occurred as a single-parameter model is constrained by very rare source populations such that, beyond a certain level, increasing disassortment would imply negative expected counts for the rarest genotypes. This boundary behaviour therefore reflects the challenge of estimating a single assortment parameter to explain tendencies across all sourc populations, but estimating individual parameters for each source population would be infeasible (though we explore this variation in the subsequent section). The model residuals show that hybrid genotypes are still generally underestimated, and pure genotypes overestimated (Table S5), indicating that disassortment is likely stronger than we are able to estimate with this constraint. The remaining five lakes showed no evidence of non-random mating (Table S4).

**Figure 2:**
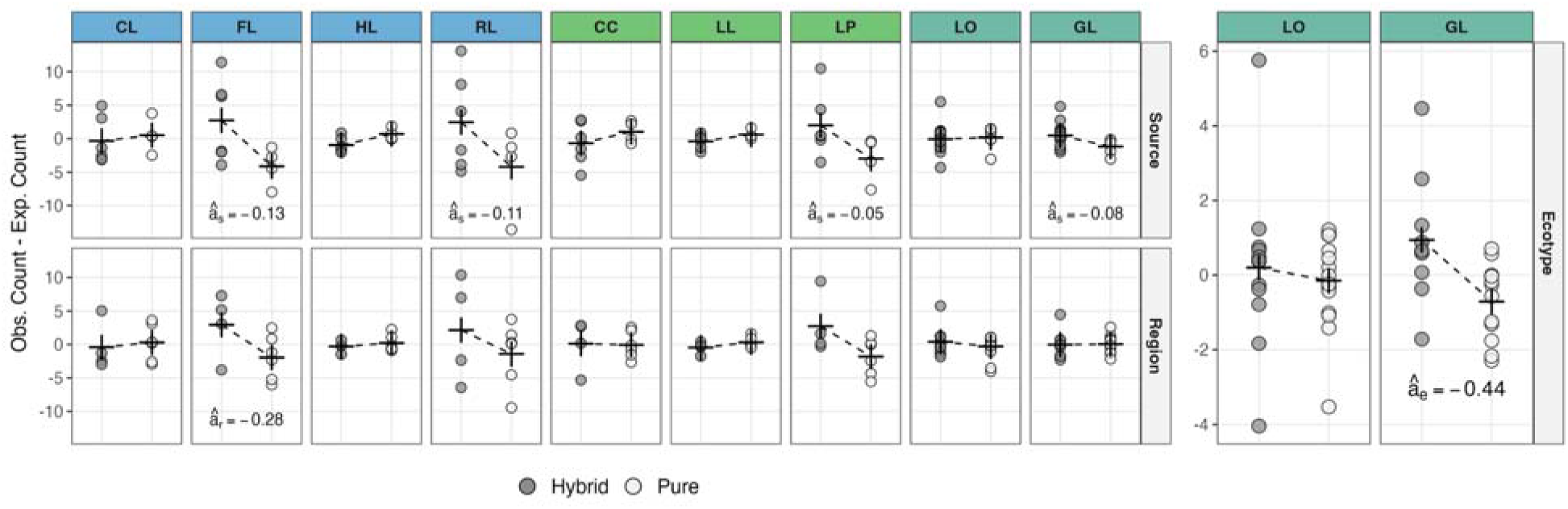
Assortment by source population, region, and ecotype in each lake: Each point represents the difference between the observed and expected counts for a particular genotype, which are grouped by whether they are pure or hybrid according to source population, region, and ecotype. For assortment by source population, expected counts are calculated directly from Hardy-Weinberg equilibrium, while for assortment by region and ecotype, expected counts are from the model accounting for assortment by source population (*â_s_*). Crossbars represent the respective means for hybrid and pure genotypes, connected by dashed lines. Assortment parameters (*â*) are shown in cases with some evidence of assortment according to Likelihood Ratio Tests, though only marginally significant in some cases. Lakes are coloured according to the ecotype of fish received (blue for limnetic, green for benthic, and teal for both).

#### Source population specific patterns

Considerable variation was evident in how individual source populations contributed to the level of disassortment in each lake (Figure 3). This variation was most consequential in Ranchero Lake (RL), where the observed pattern of disassortment seems to be driven almost entirely by behaviour of a single source population. Fish of Long Lake (LG) ancestry showed very strong disassortment (F*_i_* = −0.37), with pure LG fish being vastly under-represented; we observed just 12 pure LG fish, far fewer than the 26 expected under random mating. This single source population had an outsized influence on the overall mating dynamics in this lake because it was the most abundant, with 75% of fish (108/143) having some ancestry from LG. If we remove all fish with LG ancestry from the analysis, the estimated assortment parameter (*â_s_*) in that lake changes from −0.11 to just −0.03, much more indicative of random mating. Variation among individual source populations can be seen in the other lakes with evidence of disassortative mating, though not as heavily influenced by a single population; importantly, some evidence of disassortative mating was present in multiple lakes that did not receive LG fish as a source population (LP and GL), meaning our findings were not solely driven by tendencies of this single source population. Generally, variation was evident among source populations in the tendency for disassortment (Figure 3), with LG showing the most disassortment across all populations (mean F*_i_*= −0.14). No populations showed consistent positive assortment (i.e., mated with their own source population more than expected by chance); indeed, the highest mean F*_i_* was just 0.02 for fish from Walby Lake (WB).

**Figure 3:**
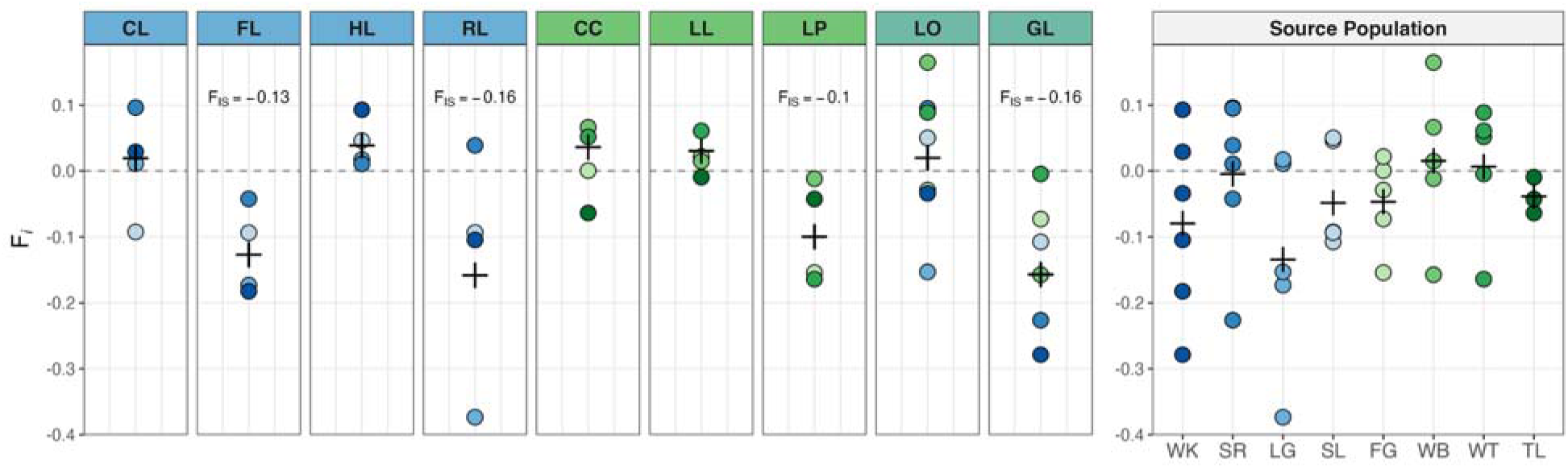
Source population specific mating patterns: Left plots show source-population-specific inbreeding coefficients (F*_i_*) for each source population in each lake. Crossbars represent the overall F_IS_ for each lake, the value of which are shown in cases with evidence of non-random assortment by population (though only marginally significant in LP). Lakes are coloured according to the ecotype of fish received (blue for limnetic, green for benthic, and teal for both). The plot on the right shows the same F*_i_* values sorted by source population, showing how th tendency for disassortment varies among source populations. Here, crossbars represent mean F*_i_* for that source population across all lakes.

#### Assortment by region

We found no conclusive evidence for non-random assortment by region (Figure 2; Table S6), though in Fred Lake (FL), we estimated strong disassortment by region ( = −0.280) even while estimating considerable disassortment by source population ( = −0.090); however the confidence intervals were wide ( 95% CI = −0.493 – 0.002), making the effect insignificant (LR = 3.78, p = 0.052, p_adj_ = 0.466). In several other cases, the magnitude of the regional assortment parameter ( ) was also quite negative ( < −0.2 for LP, RL, and LO), though the confidence intervals were always very wide (Figure S5). If we did not account for assortment by population in the same model, we would find evidence of disassortment by region in most of the lakes showing disassortment by population (Table S8), highlighting the importance of simultaneously considering both mechanisms.

#### Assortment by ecotype

Though inconclusive, we estimated very strong disassortative mating by ecotype in one of the lakes (GL) that received both ecotypes ( = −0.444), where we observed substantially more between-ecotype hybrid individuals than expected by chance (Figure 2; Table S6). However, uncertainty was high (Figure S5; 95% CI = −0.711 – 0.011), and the test was only nearly significant before multiple-test correction (LR = 3.70, p = 0.054, p_adj_ = 0.109, n = 52). Including the ecotype assortment parameter also shifted the estimated population assortment parameter in a negative direction ( = −0.146), indicating that ecotype- and population-level effects compete to explain the observed heterozygote excess. As a result, although the data are consistent with substantial disassortment in GL, they do not allow precise and confident attribution of that disassortment to ecotype versus population. No evidence of non-random mating was present in the other lake that received both ecotypes (LO: = −0.104, LR = 0.26, p_adj_ = 0.608).

### Assortment by morphological and genetic divergence

Among hybrid genotypes (according to source population), we found no conclusive evidence for non-random assortment by morphological or genetic divergence (Figure 4). Across all lake when analyzed together, the frequency of genotypes was not predicted by the degree of morphological or genetic divergence between parental populations (Figure 4; Table S9), including in a model that included just the subset of four lakes (FL, RL, LP, GL) that contained some evidence of disassortment by source population (Table S9).

**Figure 4:**
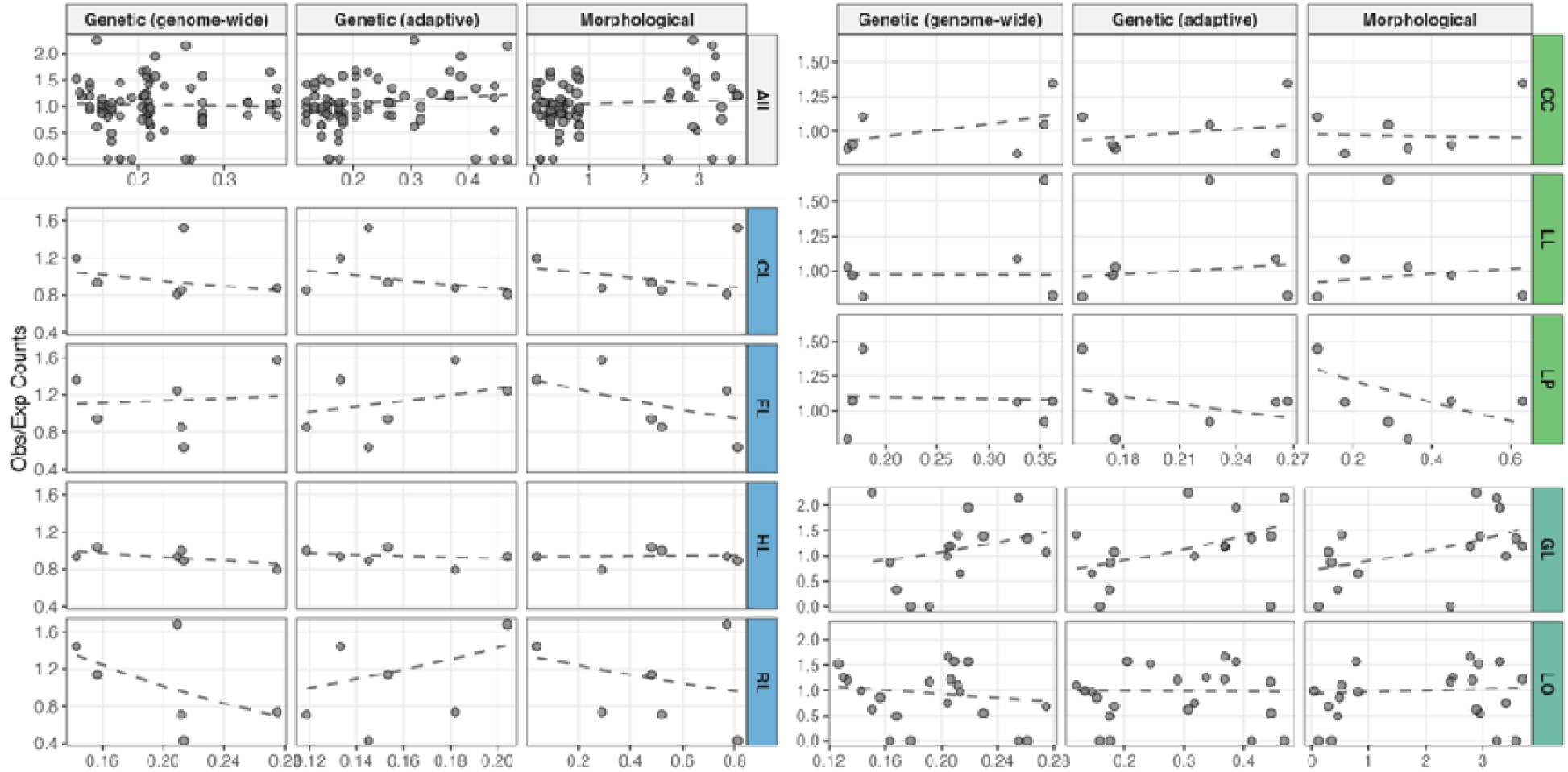
Assortment by genetic and morphological divergence: The ratio of the observed to expected counts (according to HWE) of each hybrid genotype in each lake, compared to the degree of divergence between the parental populations. Genome-wide divergence is F_ST_ across all autosomes, adaptive genetic divergence is F_ST_ across putatively adaptive windows, and morphological divergence is the difference in linear discriminant (LD) scores of morphological traits. Dashed lines represent trends from generalized linear models, none of which were significant (Table S9). Lakes are coloured according to the ecotype of fish received (blue for limnetic, green for benthic, and teal for both).

Among models for individual lakes (Figure 4; Table S9), noteworthy effects were only present in GL and RL. In GL, genotypes with greater morphological divergence (β = 0.285, SE = 0.155) and greater adaptive genetic divergence (β = 0.279, SE = 0.156) between parental populations tended to be over-represented, though both trends were only nearly significant before correcting for multiple testing (p = 0.066, p_adj_ = 0.507 and p = 0.073, p_adj_ = 0.657, respectively). Still, these results are in line with the over-representation of between-ecotype hybrid genotypes in this lake. In RL, genotypes with greater genome-wide genetic divergence tended to be under-represented (β = −0.235, SE = 0.123), although again not significant (p = 0.057, p_adj_ = 0.511). Interestingly, this trend would contradict the presence of disassortative mating in that lake.

### Selection on hybrid genotypes

Using standard length and at a given age and body condition as proxies for fitness, we found no evidence of selection in favour of hybrid genotypes in any of the lakes that showed evidence of disassortative mating (Figure S6; Figure S7; Table S10; Table S12). Because fish of LG ancestry had an outsized influence on disassortment in RL, we also compared pure LG fish to hybrid LG fish in just that lake, again finding no difference in standard length (β = −2.19, SE = 1.841, p = 0.24), or body condition (β = −0.48, SE = 0.48, p = 0.32). In GL, where between-ecotype hybrids were greatly over-represented, ecotype hybrids tended to be shorter than pure limnetic genotypes, which were the dominant genotype in the lake (although this trend was non-significant; β = −1.25, SE = 1.11, p_adj_ = 0.266), again providing no evidence of selection in favour of hybrids. Moreover, we found no differences in the proportion of hybrid fish in year one versus year two in any lake (Figure S8; Table S13), providing no evidence of differential survival between pure and hybrid fish. Although neither of these tests can rule out the possibility that differential viability or survival contributed to the observed genotype counts, the results are consistent with non-random mating being the primary mechanism.

## Discussion

Contrary to expectations from the literature, we found no evidence of assortative mating, and thus no progress towards reproductive isolation, among allopatric populations when brought into secondary contact. Instead, we found evidence of disassortative mating by population in three of our nine lakes (with weak evidence in a fourth) and some evidence of disassortative mating by ecotype in one of the two lakes that received both ecotypes. Although we cannot rule out the possibility that differential viability or survival contributed to these patterns, a lack of evidence for selection is consistent with mate choice primarily determining the observed genotype frequencies.

Assortative mating appears generally to be common (Janicke et al., 2019; Jiang et al., 2013), but it should not be entirely surprising that we failed to find any evidence in our study. Primarily, our finding likely reflects the challenge of generating reproductive isolation merely as a byproduct of divergence in allopatry, without reinforcement through secondary contact (Liou & Price, 1994). Two other studies that examined assortative mating among allopatric stickleback populations found some evidence of positive assortment (Roesti et al., 2025; Vines & Schluter, 2005), indicating that the evolution of assortative mating can vary across systems and contexts. One possible explanation for these contrasting results is that variation in the degree and direction of adaptive divergence among populations and ecotypes influences the extent of reproductive isolation (Funk et al., 2006; Nosil et al., 2009). Variation in the presence and strength of assortative mating might generally be common as several other studies of stickleback, even in sympatric and parapatric systems, have failed to find evidence of assortative mating (Jones et al., 2008; Raeymaekers et al., 2010; Räsänen et al., 2012). We suggest that future studies increasingly leverage this variation in outcomes to gain a better understanding of the determinants of progress toward reproductive isolation.

In our study, what is perhaps more surprising than the absence of assortative mating is the presence of disassortative mating in several cases. Generally, the main selective benefit of disassortative mating is inbreeding avoidance (Pusey & Wolf, 1996; Waser, 1993). This particular mechanism might seem an unlikely explanation in our case as even small bodies of water can support large populations of stickleback with high genetic diversity, where the risk of inbreeding should be low (DeFaveri & Merilä, 2015). However, mating preferences can also be influenced by evolutionary history in addition to the current context (Svensson & Gosden, 2007). For example, disassortative mating could be favoured during the process of colonizing new environments (such as stickleback colonizing lakes), when populations are small and not yet locally adapted. Then, without subsequent secondary contact to favour assortative mating, tendencies for disassortment might persist.

Although patterns of disassortative mating were evident in several replicate lakes, the cues mediating mate choice in these cases remain unknown. The frequency of hybrid genotypes was unrelated to the degree of genetic or morphological divergence between parental populations, suggesting that mate choice could be based on unmeasured traits, rather than general morphological or genetic dissimilarity. However, the tendency for disassortative mating by ecotype in G Lake (GL) suggests that, at least in certain cases, disassortment can involve traits associated with the primary axis of ecological divergence. Still, many traits do evolve largely independently of the benthic-limnetic axis, providing alternative cues. Immune-related traits are strong candidates, as immune complementarity influences mate choice across many taxa (Milinski, 2006; Penn & Potts, 1999), including stickleback (Milinski et al., 2005; Reusch et al., 2001), and the substantial variation in parasite communities and immune traits among our source populations provides a plausible basis for such mechanisms (Bolnick et al., 2026; Weber et al., 2022). Alternatively, the variation in parasite incidence among source populations provides the possibility that mate choice is mediated by condition rather than traits, where lower condition populations (such as those that were heavily parasitized at the time of introduction) might favour higher condition mates (Falk et al., 2012). Ultimately, mate choice is likely dictated by the combination and interaction of multiple traits and mechanisms (Candolin, 2003).

Finally, the variability in mating patterns across source populations and lakes reinforces the context-dependency of hybridization outcomes. First, certain source populations exhibited stronger tendencies toward disassortative mating, underscoring the notion that mate choice and, specifically the tendency for assortment, is shaped by past evolutionary history and can vary substantially among populations (Fargevieille et al., 2017; Schwartz et al., 2010; Svensson & Gosden, 2007). This finding therefore cautions against relying on inferences from just a few populations when attempting to characterize general mating patterns for an ecotype or species.

Second, despite having multiple replicates of the same set of introduced source populations, evidence of disassortative mating was present only in some replicates, demonstrating the context-dependency of mating dynamics (Gompert et al., 2017; Harrison & Larson, 2016). Ecological and environmental change can alter mating dynamics (Ålund et al., 2023; Candolin, 2019; Chunco, 2014; Ottenburghs, 2021) as exemplified in stickleback by the collapse of the benthic-limnetic species pair in Enos Lake (Taylor et al., 2006). Despite long term differentiation between sympatric ecotypes in this lake, the population gradually collapsed into a hybrid swarm (Kraak et al., 2001), plausibly due to the introduction of an invasive species altering the ecological context and impacting premating isolation and selection on hybrid phenotypes (Behm et al., 2010; Kinney et al., 2025). We suggest that similar ecological dependencies are at play in our study system given the substantial environmental and ecological divergence among our lakes (Hendry et al., 2024). More broadly, the context-dependency that we observed challenges the general validity of extending results from laboratory trials to mating dynamics in nature (Dougherty, 2020).

Assortative mating is a critical mechanism in ecological speciation, yet it does not inevitably evolve among ecologically divergent populations. Here, we use replicated whole-lake introductions to provide a rare experimental test of the expectation of assortative mating in nature, and show that it is not only absent, but that there is evidence of disassortative mating instead. These findings demonstrate that allopatric adaptive divergence alone might be insufficient to generate reproductive isolation in many cases, and additional mechanisms, such as secondary contact, might be necessary to reinforce mate choice. Further, our findings suggest a critical role of past evolutionary history and current ecological context in determining when assortative and disassortative mating are realized. Contexts generating disassortative mating pose an additional hurdle for ecological speciation to overcome on the path toward reproductive isolation, making successful cases of speciation all the more remarkable.

## Acknowledgements

Our field sites reside on the traditional and current lands of the Dena’ina Peoples, as well as private landowners, and we are grateful for the continued support of all those who allow us to work there. Fieldwork was made possible by collaboration with the Alaska Department of Fish and Game, the Salamatof Native Association, and the Kenai National Wildlife Refuge, as well as the help of many individuals who we list in the Supplementary Materials. Finally, we thank Antoine Paccard, Ariane Boisclair, and Lena Li Chun Fong of the McGill Genome Centre for collaboration with DNA extraction and genotyping.

## Author Contributions

Conceptualization: LE, DIB, CLP, APH, RDHB. Data Curation: LE. Formal Analysis: LE. Funding Acquisition: DIB, CLP, APH, RDHB. Investigation: LE. Methodology: LE. Project Administration: DIB, CLP, APH, RDHB. Supervision: DIB, CLP, APH, RDHB. Visualization: LE. Writing – original draft: LE. Writing – review and editing: LE, DIB, CLP, APH, RDHB.

## Data and Code Availability

All data and code will be archived in Figshare upon acceptance.

