## Supplementary Methods for "Disassortative mating in replicated secondary contact experiments in nature"

**Supplementary Materials**

**Supplementary Methods**

G Lake

One of the lakes that received both ecotypes is exceptional for several reasons. G Lake (GL) was originally stocked with just benthic fish in 2019, but it became apparent that the introduction failed, as no fish were caught in sampling efforts in the two years following the introduction (Hendry et al., 2024). In 2022, it was restocked with both benthic and limnetic fish, although one of the source populations (Long Lake: LG) had become depleted (probably due to invasive pike) to the point it could not be used as a source for introductions – thus, GL received seven of the eight source populations. Further, GL was stocked with a greater number of individuals than the other lakes to increase the chance of success. For those reasons, GL does not represent a true replicate of the other lake that received both benthic and limnetic fish (Loon Lake: LO) but it still provides another useful case of a lake that received both ecotypes.

Age class inference

Our analyses are designed to include only fish from the F1 generation, such that we can confidently infer mating patterns among the parental populations. Identifying these fish is made challenging by the presence of overlapping generations – as breeding can occur at one or two years of age. To discriminate between generations, we used information from both the genotyping data and measurements of standard length for each fish.

The sample of fish collected one-year post-introduction potentially contained both the F1 and parental generations, the latter representing the original fish that were transplanted. Given the rapid growth of the populations, the vast majority of fish sampled should be from the F1 generation. Although we only had standard length data for 40 of the 95 fish genotyped from each lake in this year, we were able to support this assumption by visualizing the standard lengths of fish confirmed to be from the F1 generation (Figure S1), based on having ancestry proportions of 0.5 from the genotyping data. In all lakes, F1 fish (n=306) spanned the distribution of standard lengths, save three outliers – unusually large fish that were not confirmed to be from the F1 which we deemed to be from the parental generation. Although we would miss any outliers in the additional fish without standard length data, these outliers represented a very small percentage of our sample. Though we had no means of confirming that the remaining pure fish (of similar sizes to the confirmed F1 fish) were indeed from the F1 generation, we believe this is a reasonable assumption (again due to the apparent population growth). The possibility of erroneously including fish from the parental generation would bias against our main findings of disassortative mating.

The sample of fish collected two years post-introduction (for which we have complete standard length data) potentially contained both the F1 and F2 generations, with the chance of catching any fish from parental generation being exceedingly low. We visualized the standard lengths of confirmed F2 individuals (any fish with an ancestry proportion of 0.25 from a source population) in concert with k-means clustering (k=2) of those standard lengths (Figure S2). In most lakes sampled, there was a clear bimodal distribution of standard lengths, with the confirmed F2 individuals mapping to the smaller cluster (e.g., GL in Figure S2). In these cases, we retained fish in the larger cluster (representing the second year F1 fish) and omitted any remaining confirmed F2 individuals. However, in three lakes, we appeared to sample almost exclusively the F1 generation, with very few confirmed F2 individuals and a unimodal distribution of standard lengths (e.g., LP in Figure S2). In those cases, we simply removed any confirmed F2 individuals. Even if there were additional F2 individuals that genotypically resembled F1s, those individuals would necessarily be the result of assortative mating in the first generation to produce pure F1 fish; again, implying that any potential bias should oppose our main finding of disassortative mating.

Models of assortative mating

To quantify the degree of non-random mating within each lake, we estimated an assortment parameter ($a_{s}$) using a maximum likelihood approach. This parameter is analogous to Wright’s fixation index or inbreeding coefficient (F_IS_) but is used here to test for assortative or disassortative mating. Specifically, we modified the Hardy-Weinberg expectations for genotype frequencies of homozygotes and heterozygotes to incorporate our assortment parameter.

Homozygotes: $P_{ii}=p_{i}^{2}+a_{s}p_{i}(1-p_{i})$

Heterozygotes: $P_{ij}=2p_{i}p_{j}(1-a_{s})$

Here, $a_{s}=0$ would represent random mating, $a_{s}=1$ would represent perfect assortative mating, and $a_{s}=-1$ would represent perfect disassortative mating. We estimated $\hat{a}_{s}$ by maximizing the log-likelihood of the observed genotype counts across the state space of all $k$ populations present in each lake. The log-likelihood function ($l$) is defined as:

$$l\left( a_{s} | n \right)=\sum_{g=1}^{G} n_{g}ln(P_{g}\left( a_{s} \right))$$

Here, $n_{g}$ is the observed count of genotype $g$, and $P_{g}\left( a_{s} \right)$ is the expected probability of that genotype given $a_{s}$. We restricted $\hat{a}_{s}$ to be between -0.999 and 0.999 to ensure numerical stability and probabilistic validity. Optimization was performed using the Nelder-Mead algorithm via the optim function in R and we extracted confidence intervals using profile likelihood estimation. We then tested for statistical significance using a Likelihood Ratio Test (LRT) against a model of random mating ($a_{s}=0$).

To test for assortment by region and ecotype, we extended our model to include a hierarchical structure. This model estimates two parameters simultaneously: $a_{s}$ for assortment by source population and either $a_{r}$ for assortment by region or $a_{e}$ for assortment by ecotype. For example, for assortment by ecotype, the probabilities of each genotype are defined as:

Homozygotes: ${P'}_{ii}=\left[ p_{i}^{2}+a_{s}p_{i}\left( 1-p_{i} \right) \right]\times(1+a_{e})$

Heterozygotes (same ecotype): ${P'}_{ij}=[2p_{i}p_{j}\left( 1-a_{s} \right)]\times(1+a_{e})$

Heterozygotes (different ecotype): ${P'}_{ij}=2p_{i}p_{j}\left( 1-a_{s} \right)$

Because the addition of the second parameter ($a_{s}$) shifts the probability space, we performed a global normalization to ensure the final probabilities summed to 1. We estimated both parameters by maximizing the joint log-likelihood across the observed genotype counts. Optimization was performed using Nelder-Mead algorithm via the optim function in R and we extracted confidence intervals using profile likelihood estimation. To test for significance of this model, we again used a Likelihood Ratio Test (LRT), comparing against a model with assortment by source population ($a_{s}$) but no assortment by region ($a_{r}=0$) or ecotype ($a_{e}=0$).

Population genetic metrics

We also used traditional population genetic tests to support results from the above assortment tests. Specifically, we used exact tests of Hardy-Weinberg equilibrium (HWE), implemented through the *pegas* R package (v.1.4), which test if the genotype frequencies (in the same sense as above) differ significantly from HWE (Guo & Thompson, 1992; Paradis, 2010). Concurrently, we computed observed heterozygosity (H_o_), expected heterozygosity (H_e_), and inbreeding coefficients (F_IS_) within each lake using the *hierfstat* package (v.0.5-11; Goudet, 2005).

Testing for deviations from random mating using exact tests of Hardy-Weinberg equilibrium (HWE) yielded similar conclusions (Table S7) to our assortment parameter estimation. Although $\hat{a}_{s}$ and F_IS_ are conceptually similar, numerical differences arose in their estimates, with estimates of $\hat{a}_{s}$ being generally lower in magnitude. The approaches also differed slightly in statistical significance outcomes, which is to be expected as exact HWE tests assess any deviations from HWE whereas our approach directly tests for assortative mating. Still, results from the HWE tests support non-random mating in FL and RL and suggest weak evidence for non-random mating in GL and LP, with F_IS_ values indicating disassortment in these cases.

Genetic divergence

To quantify genetic divergence, we estimated two different metrics of pairwise F_ST_ between source populations: one across the entire genome (genome-wide divergence) and one across just putatively adaptive windows in the genome (adaptive divergence). For genome-wide genetic divergence, we estimated F_ST_ across just the autosomes, removing the sex chromosomes, mitochondrial sequences, and other scaffolds. For adaptive genetic divergence, we identified regions putatively associated with adaptation along the benthic-limnetic divergence. Although we recognize that this contrast captures only one specific axis of adaptation, it should parallel the major axis of morphological and ecological variation among these populations. To identify putatively adaptive regions, we calculated F_ST_ in 10kb windows across the autosomes, computed the mean F_ST_ of between-ecotype population comparisons and within-ecotype population comparisons, computed the difference between those two metrics for each window, and selected the 1% most divergent windows (Figure S3). We then computed pairwise F_ST_ across those putatively adaptive windows. The validity of these adaptive F_ST_ metrics were supported through positive correlation with morphological divergence and clustering of ecotypes in a neighbour-joining tree (Figure S4).

To calculate F_ST_ metrics, we accessed raw pool-seq reads of each source population from Weber et al. (2022), trimmed them using *fastp* (v.0.24.0), and removed reads with Phred scores less than 20 or lengths shorter than 50bp, as well as adapter sequences and polyG tails. We performed quality control on both the raw and trimmed reads using *fastQC* (v.0.12.1) and *MultiQC* (v.1.31). The trimmed reads were then aligned to the University of Georgia *G. aculeatus* reference genome v.5 (Nath et al., 2021) using *BWA* (v.0.7.18), filtering out reads with a mapping score less than 20. The resulting bam files were sorted and indexed with *Samtools* (v.1.20), and PCR duplicates were marked and removed using *Picard* (v.3.1.0). We identified and masked indel regions (5bp above and below each indel) using *PoPoolation2* (Kofler et al., 2011), before converting to a sync file and filtering out positions with a base quality less than 20.

We computed all F_ST_ metrics using *grenedalf* v.0.6.3 (Czech et al., 2024). In each case, we filtered out positions with a minimum read count less than 2, a minimum depth less than 4, or a maximum depth greater than 200, and averaged across all remaining positions, either across the genome (for genome-wide divergence) or across putatively adaptive windows (for adaptive divergence). Resulting F_ST_ estimates for genome-wide and adaptive divergence are in Table S3.

Extended Acknowledgements

We thank the fieldwork crews that conducted the transplant and collected fish in 2019, 2020, 2021, 2022, and 2024, including Ismail Ameen, Luis Baertschi, Tina Barbasch, Steven Bezdecny, Chelsea Bishop, Chad Brock, Sheila Christen, Mariane Daneau-Lamoureux, Alison Derry, Victor Frankel, Gregor Fussmann, Alex Gouvin Moffat, Stéphanie Guernon, Alexis Heckley, Rachel Kramp, Kelly Ireland, Matt Josephson, Åsa Lind, Zuyao Liu, Kathryn Milligan-McClellan, Louis Moisan, Anya Mueller, Kevin Neumann, Michelle Packer, Arshad Padhiar, Sarah Pasqualetti, Allegra Pearce, Christopher Peterson, Hillary Poore, Maria Rodgers, Allison Roth, Andrea Roth, Rogini Runghen, Sarah Sanderson, Trey Sasser, Natalie Steinel, Hiranya Sudasinghe, Richard Benjamin Sulser, Saraswathy Vaidyanathan, Matthew Walsh, Jesse Weber, and Anika Wohlleben.

**Supplementary Tables**

**Table S1:** Summary of source and recipient lakes. For source lakes, ecotype describes the ecotype of the population according to morphology; for recipient lakes, ecotype describes the ecotype of populations introduced to that lake.

| **Abbr.** | **Name** | **Type** | **Ecotype** | **Region** | **Coordinates** |
| --- | --- | --- | --- | --- | --- |
| FG | Finger Lake | Source | Benthic | Mat-Su | 61.606 N, 149.279 W |
| TL | Tern Lake | Source | Benthic | Kenai | 60.533 N, 149.550 W |
| WB | Walby Lake | Source | Benthic | Mat-Su | 61.620 N, 149.213 W |
| WT | Watson Lake | Source | Benthic | Kenai | 60.539 N, 150.467 W |
| LG | Long Lake | Source | Limnetic | Mat-Su | 61.578 N, 149.764 W |
| SL | Spirit Lake | Source | Limnetic | Kenai | 60.593 N, 150.986 W |
| SR | South Rolly Lake | Source | Limnetic | Mat-Su | 61.667 N, 150.138 W |
| WK | Wik Lake | Source | Limnetic | Kenai | 60.720 N, 151.250 W |
| CC | CC Lake | Recipient | Benthic | Kenai | 60.422 N, 151.195 W |
| LL | Leisure Lake | Recipient | Benthic | Kenai | 60.415 N, 151.210 W |
| LP | Leisure Pond | Recipient | Benthic | Kenai | 60.419 N, 151.207 W |
| CL | Crystal Lake | Recipient | Limnetic | Kenai | 60.424 N, 151.194 W |
| FL | Fred’s Lake | Recipient | Limnetic | Kenai | 60.423 N, 151.200 W |
| HL | Hope Lake | Recipient | Limnetic | Kenai | 60.422 N, 151.188 W |
| RL | Ranchero Lake | Recipient | Limnetic | Kenai | 60.423 N, 151.183 W |
| GL | G Lake | Recipient | Both | Kenai | 60.430 N, 151.177 W |
| LO | Loon Lake | Recipient | Both | Kenai | 60.520 N, 151.051 W |

**Table S2**: Stocking proportions. The number of fish from each source population (columns) introduced to each recipient lake (rows) after accounting for mortalities during translocation.

|  | **LG** | **SL** | **SR** | **WK** | **FG** | **TL** | **WB** | **WT** | **Total** |
| --- | --- | --- | --- | --- | --- | --- | --- | --- | --- |
| **CL** | 366 | 393 | 388 | 270 | 0 | 0 | 0 | 0 | 1417 |
| **FL** | 99 | 100 | 90 | 97 | 0 | 0 | 0 | 0 | 386 |
| **HL** | 430 | 452 | 400 | 287 | 0 | 0 | 0 | 0 | 1569 |
| **RL** | 193 | 199 | 180 | 151 | 0 | 0 | 0 | 0 | 723 |
| **CC** | 0 | 0 | 0 | 0 | 202 | 179 | 198 | 202 | 781 |
| **LL** | 0 | 0 | 0 | 0 | 359 | 243 | 391 | 399 | 1392 |
| **LP** | 0 | 0 | 0 | 0 | 94 | 95 | 100 | 104 | 393 |
| **LO** | 245 | 295 | 242 | 132 | 288 | 130 | 282 | 287 | 1901 |
| **GL** | 0 | 500 | 500 | 495 | 500 | 500 | 500 | 500 | 3495 |

**Table S3:** Morphological and genetic divergence among source populations. Morphological is based on the absolute difference between linear discriminant (LD) scores quantifying morphological variation among populations, described in Hendry et al. (2024). Genome-wide genetic divergence is F_ST_ across all autosomes. Adaptive genetic divergence in F_ST_ in putatively adaptive windows in the genome identified by contrasting the benthic and limnetic populations (Figure S3).

| **Pop** | **Pop** | **Ecotype** | **Morphological** | **Genetic (genome-wide)** | **Genetic (adaptive)** |
| --- | --- | --- | --- | --- | --- |
| FG | LG | Inter | 2.81 | 0.133 | 0.289 |
| FG | SL | Inter | 2.77 | 0.205 | 0.368 |
| FG | SR | Inter | 3.29 | 0.219 | 0.387 |
| FG | TL | Intra | 0.29 | 0.354 | 0.226 |
| FG | WB | Intra | 0.34 | 0.163 | 0.175 |
| FG | WK | Inter | 3.58 | 0.261 | 0.413 |
| FG | WT | Intra | 0.11 | 0.178 | 0.158 |
| LG | SL | Intra | 0.04 | 0.143 | 0.133 |
| LG | SR | Intra | 0.48 | 0.157 | 0.153 |
| LG | TL | Inter | 3.10 | 0.316 | 0.428 |
| LG | WB | Inter | 2.47 | 0.131 | 0.337 |
| LG | WK | Intra | 0.77 | 0.209 | 0.204 |
| LG | WT | Inter | 2.92 | 0.127 | 0.243 |
| SL | SR | Intra | 0.52 | 0.212 | 0.119 |
| SL | TL | Inter | 3.06 | 0.349 | 0.504 |
| SL | WB | Inter | 2.43 | 0.192 | 0.445 |
| SL | WK | Intra | 0.81 | 0.214 | 0.145 |
| SL | WT | Inter | 2.88 | 0.151 | 0.306 |
| SR | TL | Inter | 3.58 | 0.379 | 0.496 |
| SR | WB | Inter | 2.95 | 0.230 | 0.446 |
| SR | WK | Intra | 0.29 | 0.275 | 0.182 |
| SR | WT | Inter | 3.40 | 0.204 | 0.317 |
| TL | WB | Intra | 0.63 | 0.362 | 0.267 |
| TL | WK | Inter | 3.87 | 0.404 | 0.552 |
| TL | WT | Intra | 0.18 | 0.328 | 0.261 |
| WB | WK | Inter | 3.24 | 0.255 | 0.468 |
| WB | WT | Intra | 0.45 | 0.168 | 0.174 |
| WK | WT | Inter | 3.69 | 0.207 | 0.367 |

**Table S4:** Models of assortative mating by source population. The estimated assortment parameter ($\hat{a}$) for assortment by source population with 95% confidence intervals, as well as the log likelihood of that model and a likelihood ratio compared to a model of random mating.

|  | **Lake** | ${\hat{\boldsymbol{a}}}_{\boldsymbol{s}}$ | **95% CI** | **logLik** | **LR** | **p-value** | **p_adj_** | **n** |
| --- | --- | --- | --- | --- | --- | --- | --- | --- |
| Limnetic | CL | 0.004 | -0.069, 0.098 | -370.65 | 0.009 | 0.923 | 0.961 | 181 |
|  | FL | -0.128 | -0.183, -0.053 | -365.08 | 9.867 | 0.002 | 0.015 | 182 |
|  | HL | 0.040 | -0.051, 0.143 | -311.19 | 0.672 | 0.412 | 0.742 | 145 |
|  | RL | -0.109 | -0.109, -0.039 | -281.38 | 7.409 | 0.006 | 0.029 | 143 |
| Benthic | CC | 0.002 | -0.067, 0.106 | -360.50 | 0.002 | 0.961 | 0.961 | 184 |
|  | LL | 0.025 | -0.013, 0.145 | -252.35 | 0.205 | 0.651 | 0.837 | 159 |
|  | LP | -0.045 | -0.045, -0.003 | -324.30 | 4.123 | 0.042 | 0.095 | 184 |
| Both | LO | 0.023 | -0.034, 0.122 | -306.34 | 0.320 | 0.572 | 0.837 | 107 |
|  | GL | -0.083 | -0.083, -0.026 | -141.04 | 5.758 | 0.016 | 0.049 | 52 |

**Table S5:** Residuals from models of assortment by source population for the three lakes where the confidence intervals are truncated. In these cases, the models are likely underestimating the magnitude of disassortment due to mathematical constraints, as evidenced by the generally positive residuals of hybrid genotypes and negative residuals of pure genotype.

| **Lake** | **Class** | **Genotype** | **Observed** | **Expected** | **Residual** |
| --- | --- | --- | --- | --- | --- |
| RL | Hybrid | LGSL | 43 | 32.56 | 10.44 |
|  |  | LGSR | 33 | 31.63 | 1.37 |
|  |  | LGWK | 20 | 13.02 | 6.98 |
|  |  | SLSR | 12 | 18.45 | -6.45 |
|  |  | SLWK | 3 | 7.60 | -4.60 |
|  |  | SRWK | 5 | 7.38 | -2.38 |
|  |  | **Mean** | - | - | **0.893** |
|  | Pure | LGLG | 12 | 21.40 | -9.40 |
|  |  | SLSL | 6 | 5.70 | 0.30 |
|  |  | SRSR | 9 | 5.27 | 3.73 |
|  |  | WKWK | 0 | 0.00 | 0.00 |
|  |  | **Mean** | - | - | **-1.34** |
| LP | Hybrid | FGTL | 2 | 2.27 | -0.27 |
|  |  | FGWB | 14 | 18.32 | -4.32 |
|  |  | FGWT | 34 | 24.57 | 9.43 |
|  |  | TLWB | 6 | 5.86 | 0.14 |
|  |  | TLWT | 8 | 7.86 | 0.14 |
|  |  | WBWT | 65 | 63.40 | 1.60 |
|  |  | **Mean** | - | - | **1.12** |
|  | Pure | FGFG | 0 | 2.42 | -2.42 |
|  |  | TLTL | 0 | 0.00 | 0.00 |
|  |  | WBWB | 22 | 20.71 | 1.29 |
|  |  | WTWT | 33 | 38.58 | -5.58 |
|  |  | **Mean** | - | - | **-1.68** |
| GL | Hybrid | FGSL | 1 | 0.92 | 0.08 |
|  |  | FGSR | 3 | 1.67 | 1.33 |
|  |  | FGWB | 1 | 1.25 | -0.25 |
|  |  | FGWK | 3 | 2.42 | 0.58 |
|  |  | FGWT | 0 | 1.75 | -1.75 |
|  |  | SLSR | 3 | 2.29 | 0.71 |
|  |  | SLWB | 0 | 1.72 | -1.72 |
|  |  | SLWK | 2 | 3.32 | -1.32 |
|  |  | SLWT | 5 | 2.41 | 2.59 |
|  |  | SRWB | 4 | 3.12 | 0.88 |
|  |  | SRWK | 6 | 6.04 | -0.04 |
|  |  | SRWT | 4 | 4.37 | -0.37 |
|  |  | WBWK | 9 | 4.53 | 4.47 |
|  |  | WBWT | 1 | 3.28 | -2.28 |
|  |  | WKWT | 7 | 6.34 | 0.66 |
|  |  | **Mean** | - | - | **0.238** |
|  | Pure | FGFG | 0 | 0.00 | 0.00 |
|  |  | SLSL | 0 | 0.17 | -0.17 |
|  |  | SRSR | 0 | 1.25 | -1.25 |
|  |  | WBWB | 0 | 0.55 | -0.55 |
|  |  | WKWK | 1 | 3.17 | -2.17 |
|  |  | WTWT | 2 | 1.42 | 0.58 |
|  |  | **Mean** | - | - | **-0.593** |

**Table S6:** Models of assortment by region and ecotype. The estimated assortment parameter for region or ecotype and the estimated source population assortment parameter, both with confidence intervals, as well as the log likelihood of that model and a likelihood ratio compared to a model of including just assortment by source population.

|  |  | **Lake** | $\hat{\boldsymbol{a}}$ | $\hat{\boldsymbol{a}}$**: 95% CI** | ${\hat{\boldsymbol{a}}}_{\boldsymbol{s}}$ | ${\hat{\boldsymbol{a}}}_{\boldsymbol{s}}$**: 95% CI** | **logLik** | **LR** | **p-value** | **p_adj_** | **n** |
| --- | --- | --- | --- | --- | --- | --- | --- | --- | --- | --- | --- |
| Region | Limnetic | CL | 0.052 | -0.253, 0.471 | -0.002 | -0.077, 0.106 | -370.61 | 0.09 | 0.768 | 0.935 | 181 |
|  |  | FL | -0.280 | -0.493, 0.002 | -0.090 | -0.167, 0.021 | -363.19 | 3.78 | 0.052 | 0.466 | 182 |
|  |  | HL | 0.052 | -0.299, 0.552 | 0.031 | -0.07, 0.162 | -311.16 | 0.06 | 0.803 | 0.935 | 145 |
|  |  | RL | -0.224 | -0.477, 0.093 | -0.102 | -0.174, 0.023 | -280.34 | 2.09 | 0.149 | 0.669 | 143 |
|  | Benthic | CC | -0.016 | -0.354, 0.438 | 0.006 | -0.098, 0.163 | -360.50 | 0.01 | 0.935 | 0.935 | 184 |
|  |  | LL | 0.075 | -0.274, 0.564 | 0.012 | -0.09, 0.158 | -252.28 | 0.14 | 0.711 | 0.935 | 159 |
|  |  | LP | -0.038 | -0.384, 0.305 | -0.150 | -0.224, 0.019 | -322.37 | 0.03 | 0.852 | 0.935 | 184 |
|  | Both | LO | -0.208 | -0.501, 0.22 | 0.051 | -0.035, 0.188 | -305.80 | 1.09 | 0.297 | 0.890 | 107 |
|  |  | GL | 0.189 | -0.318, 1.08 | -0.173 | -0.239, -0.073 | -139.43 | 0.37 | 0.544 | 0.935 | 52 |
| Eco | Both | LO | -0.104 | -0.414, 0.361 | 0.033 | -0.038, 0.15 | -306.21 | 0.26 | 0.608 | 0.608 | 107 |
|  |  | GL | -0.444 | -0.711, 0.011 | -0.146 | -0.231, 0.011 | -137.76 | 3.70 | 0.054 | 0.109 | 52 |

**Table S7**: Traditional population genetics tests for non-random mating. The observed (H_o_) and expected (H_e_) proportions of hybrid heterozygotes and the corresponding inbreeding coefficient (F_IS_) in each lake, as well as results from exact Hardy-Weinberg equilibrium tests, adjusted for multiple testing. For assortment by region and ecotype, this approach does simultaneously account for assortment by source population, which is why we find evidence of assortment by region in the lakes with assortment by source population.

|  | **Lake** | **Admixture Metrics** | | | **HWE Tests** | | | |
| --- | --- | --- | --- | --- | --- | --- | --- | --- |
|  |  | **H_o_** | **H_e_** | **F_IS_** | **X^2^** | **df** | **p-value** | **p_adj_** |
| Source Population | CL | 0.680 | 0.693 | 0.019 | 5.80 | 6 | 0.424 | 0.763 |
|  | FL | 0.786 | 0.697 | -0.127 | 18.85 | 6 | 0.003 | 0.013 |
|  | HL | 0.683 | 0.710 | 0.039 | 1.59 | 6 | 0.936 | 0.936 |
|  | RL | 0.811 | 0.700 | -0.159 | 25.83 | 6 | 0.001 | 0.005 |
|  | CC | 0.647 | 0.671 | 0.036 | 3.57 | 6 | 0.788 | 0.936 |
|  | LL | 0.541 | 0.558 | 0.030 | 1.16 | 6 | 0.866 | 0.936 |
|  | LP | 0.701 | 0.637 | -0.100 | 10.99 | 6 | 0.079 | 0.212 |
|  | LO | 0.785 | 0.801 | 0.020 | 16.53 | 21 | 0.651 | 0.936 |
|  | GL | 0.942 | 0.813 | -0.159 | 22.91 | 15 | 0.094 | 0.212 |
| Region | CL | 0.381 | 0.393 | 0.031 | 0.15 | 1 | 0.702 | 0.872 |
|  | FL | 0.577 | 0.455 | -0.267 | 13.24 | 1 | <0.001 | 0.001 |
|  | HL | 0.448 | 0.477 | 0.061 | 0.48 | 1 | 0.487 | 0.845 |
|  | RL | 0.559 | 0.452 | -0.239 | 8.36 | 1 | 0.005 | 0.023 |
|  | CC | 0.500 | 0.500 | 0.001 | 0.001 | 1 | 1.000 | 1.000 |
|  | LL | 0.346 | 0.368 | 0.061 | 0.53 | 1 | 0.510 | 0.845 |
|  | LP | 0.582 | 0.501 | -0.161 | 4.94 | 1 | 0.034 | 0.102 |
|  | LO | 0.533 | 0.502 | -0.061 | 0.46 | 1 | 0.563 | 0.845 |
|  | GL | 0.519 | 0.489 | -0.061 | 0.26 | 1 | 0.775 | 0.872 |
| Eco | LO | 0.421 | 0.410 | -0.027 | 0.11 | 1 | 0.824 | 0.824 |
|  | GL | 0.692 | 0.489 | -0.410 | 9.09 | 1 | 0.005 | 0.009 |

**Table S8:** Testing for assortment by region and ecotype without considering assortment by source population.

|  | **Lake** | $\hat{\boldsymbol{a}}$ | **95% CI** | **logLik** | **LR** | **p-value** | **p_adj_** | **n** |
| --- | --- | --- | --- | --- | --- | --- | --- | --- |
| Region | CL | 0.028 | -0.111, 0.18 | -162.501 | 0.143 | 0.705 | 0.793 | 181 |
|  | FL | -0.270 | -0.381, -0.135 | -155.569 | 14.144 | <0.001 | 0.002 | 182 |
|  | HL | 0.058 | -0.104, 0.22 | -148.598 | 0.479 | 0.489 | 0.748 | 145 |
|  | RL | -0.242 | -0.369, -0.087 | -123.944 | 8.879 | 0.003 | 0.013 | 143 |
|  | CC | -0.002 | -0.146, 0.142 | -190.960 | 0.001 | 0.98 | 0.98 | 184 |
|  | LL | 0.057 | -0.092, 0.223 | -137.644 | 0.513 | 0.474 | 0.748 | 159 |
|  | LP | -0.164 | -0.303, -0.02 | -178.292 | 4.966 | 0.026 | 0.078 | 184 |
|  | LO | -0.065 | -0.251, 0.123 | -108.595 | 0.458 | 0.498 | 0.748 | 107 |
|  | GL | -0.071 | -0.326, 0.2 | -51.677 | 0.260 | 0.61 | 0.785 | 52 |
| Eco | LO | -0.032 | -0.2, 0.163 | -96.652 | 0.109 | 0.741 | 0.741 | 107 |
|  | GL | -0.418 | -0.61, -0.162 | -41.094 | 9.609 | 0.002 | 0.004 | 52 |

**Table S9**: Generalized linear models of assortment by morphological and genetic divergence. Mixed models with lake as a random effect were used to model across all lakes, and across the four lakes (Subset: FL, RL, LP, GL) with some evidence of disassortment. Generalized linear models were used for individual lakes and p-values were adjusted for multiple testing.

| **Lake** | **Divergence** | **Effect** | **SE** | **T-value** | **p-value** | **p_adj_** |
| --- | --- | --- | --- | --- | --- | --- |
| All | Genetic (genome-wide) | -0.235 | 0.684 | -0.344 | 0.731 | 0.731 |
| All | Genetic (adaptive) | 0.589 | 0.523 | 1.126 | 0.260 | 0.260 |
| All | Morphological | 0.027 | 0.042 | 0.635 | 0.526 | 0.526 |
| Subset | Genetic (genome-wide) | -0.439 | 1.025 | -0.429 | 0.668 | 0.668 |
| Subset | Genetic (adaptive) | 1.043 | 0.667 | 1.564 | 0.118 | 0.860 |
| Subset | Morphological | 0.041 | 0.061 | 0.678 | 0.498 | 0.498 |
| CL | Genetic (genome-wide) | -0.074 | 0.109 | -0.680 | 0.497 | 0.912 |
| CL | Genetic (adaptive) | -0.080 | 0.116 | -0.685 | 0.494 | 0.948 |
| CL | Morphological | -0.081 | 0.112 | -0.721 | 0.471 | 0.848 |
| FL | Genetic (genome-wide) | 0.027 | 0.096 | 0.280 | 0.78 | 0.996 |
| FL | Genetic (adaptive) | 0.086 | 0.090 | 0.948 | 0.343 | 0.948 |
| FL | Morphological | -0.138 | 0.087 | -1.586 | 0.113 | 0.507 |
| HL | Genetic (genome-wide) | -0.055 | 0.118 | -0.468 | 0.64 | 0.96 |
| HL | Genetic (adaptive) | -0.025 | 0.109 | -0.226 | 0.821 | 0.948 |
| HL | Morphological | 0.007 | 0.103 | 0.065 | 0.948 | 0.948 |
| RL | Genetic (genome-wide) | -0.235 | 0.123 | -1.905 | 0.057 | 0.511 |
| RL | Genetic (adaptive) | 0.141 | 0.109 | 1.297 | 0.195 | 0.875 |
| RL | Morphological | -0.122 | 0.101 | -1.210 | 0.226 | 0.514 |
| CC | Genetic (genome-wide) | 0.087 | 0.127 | 0.684 | 0.494 | 0.912 |
| CC | Genetic (adaptive) | 0.045 | 0.124 | 0.366 | 0.714 | 0.948 |
| CC | Morphological | -0.008 | 0.112 | -0.074 | 0.941 | 0.948 |
| LL | Genetic (genome-wide) | -0.001 | 0.275 | -0.004 | 0.996 | 0.996 |
| LL | Genetic (adaptive) | 0.039 | 0.261 | 0.151 | 0.88 | 0.948 |
| LL | Morphological | 0.038 | 0.172 | 0.219 | 0.827 | 0.948 |
| LP | Genetic (genome-wide) | -0.011 | 0.147 | -0.075 | 0.941 | 0.996 |
| LP | Genetic (adaptive) | -0.083 | 0.145 | -0.574 | 0.566 | 0.948 |
| LP | Morphological | -0.130 | 0.108 | -1.204 | 0.229 | 0.514 |
| LO | Genetic (genome-wide) | -0.089 | 0.135 | -0.664 | 0.507 | 0.912 |
| LO | Genetic (adaptive) | -0.008 | 0.130 | -0.065 | 0.948 | 0.948 |
| LO | Morphological | 0.043 | 0.111 | 0.387 | 0.699 | 0.948 |
| GL | Genetic (genome-wide) | 0.151 | 0.142 | 1.062 | 0.288 | 0.912 |
| GL | Genetic (adaptive) | 0.279 | 0.156 | 1.793 | 0.073 | 0.657 |
| GL | Morphological | 0.285 | 0.155 | 1.837 | 0.066 | 0.507 |

**Table S10**: Selection on hybrid genotypes. Linear models of standard length as a function of whether the genotype is pure or hybrid by source population, as well as age (as our dataset includes fish of age 1 and 2). All p-values were adjusted for multiple testing.

| **Lake** | **Term** | **Estimate** | **SE** | **T-statistic** | **p-value** | **p_adj_** |
| --- | --- | --- | --- | --- | --- | --- |
| CL | Genotype (pure) | 0.470 | 0.835 | 0.563 | 0.575 | 0.715 |
| CL | Age (2) | 8.790 | 0.874 | 10.060 | <0.001 | <0.001 |
| FL | Genotype (pure) | 0.371 | 1.120 | 0.331 | 0.741 | 0.741 |
| FL | Age (2) | 10.954 | 1.023 | 10.706 | <0.001 | <0.001 |
| HL | Genotype (pure) | 0.613 | 1.109 | 0.552 | 0.582 | 0.715 |
| HL | Age (2) | 8.179 | 1.109 | 7.373 | <0.001 | <0.001 |
| RL | Genotype (pure) | 1.059 | 1.376 | 0.769 | 0.444 | 0.715 |
| RL | Age (2) | 8.180 | 1.100 | 7.436 | <0.001 | <0.001 |
| CC | Genotype (pure) | -1.044 | 0.708 | -1.475 | 0.143 | 0.429 |
| CC | Age (2) | -0.425 | 0.745 | -0.570 | 0.57 | 0.57 |
| LL | Genotype (pure) | -2.904 | 0.935 | -3.108 | 0.002 | 0.022 |
| LL | Age (2) | 12.463 | 0.981 | 12.698 | <0.001 | <0.001 |
| LP | Genotype (pure) | -1.866 | 0.880 | -2.121 | 0.036 | 0.162 |
| LP | Age (2) | 7.419 | 0.865 | 8.575 | <0.001 | <0.001 |
| LO | Genotype (pure) | -1.491 | 1.607 | -0.928 | 0.358 | 0.715 |
| LO | Age (2) | 12.429 | 1.447 | 8.592 | <0.001 | <0.001 |
| GL | Genotype (pure) | 0.984 | 2.064 | 0.476 | 0.636 | 0.715 |

**Table S11**: Selection on hybrid genotypes by ecotype. Linear models of standard length as a function of whether that genotype is hybrid or pure with respect to ecotype, as well as age.

| **Lake** | **Term** | **Estimate** | **SE** | **T-statistic** | **p-value** | **p_adj_** |
| --- | --- | --- | --- | --- | --- | --- |
| LO | Pure Benthic | 2.251 | 2.633 | 0.855 | 0.397 | 0.397 |
| LO | Pure Limnetic | -2.529 | 1.242 | -2.036 | 0.048 | 0.095 |
| LO | Age (2) | 11.624 | 1.444 | 8.049 | <0.001 | <0.001 |
| GL | Pure Benthic | -3.314 | 1.754 | -1.890 | 0.065 | 0.129 |
| GL | Pure Limnetic | 1.247 | 1.109 | 1.124 | 0.266 | 0.266 |

**Table S12**: Selection on hybrid genotypes. Linear models of body condition as a function of whether the genotype is pure or hybrid by source population. All p-values were adjusted for multiple testing.

| **Lake** | **Term** | **Estimate** | **SE** | **T-statistic** | **p-value** | **p_adj_** |
| --- | --- | --- | --- | --- | --- | --- |
| CL | Genotype (pure) | -0.132 | 0.224 | -0.591 | 0.556 | 0.556 |
| FL | Genotype (pure) | -0.207 | 0.247 | -0.839 | 0.404 | 0.556 |
| HL | Genotype (pure) | -0.183 | 0.279 | -0.657 | 0.514 | 0.556 |
| RL | Genotype (pure) | -0.230 | 0.370 | -0.622 | 0.537 | 0.556 |
| CC | Genotype (pure) | 0.334 | 0.219 | 1.525 | 0.131 | 0.458 |
| LL | Genotype (pure) | 0.267 | 0.250 | 1.069 | 0.289 | 0.556 |
| LP | Genotype (pure) | 0.383 | 0.234 | 1.639 | 0.105 | 0.458 |
| LO | Genotype (pure) | 1.084 | 0.737 | 1.472 | 0.172 | 0.458 |

**Table S13**: Selection on hybrid genotypes. Fisher’s exact tests of differences in the proportion of hybrid individuals between the first and second year of sampling in each lake. We could not do this test in G Lake (GL) as we only sampled that lake in year two.

| **Lake** | **Δ proportion** | **p-value** | **p_adj_** |
| --- | --- | --- | --- |
| CL | -0.098 | 0.202 | 0.538 |
| FL | -0.092 | 0.143 | 0.538 |
| HL | -0.126 | 0.136 | 0.538 |
| RL | 0.065 | 0.498 | 0.995 |
| CC | -0.019 | 0.877 | 1 |
| LL | -0.016 | 0.872 | 1 |
| LP | 0.035 | 0.632 | 1 |
| LO | 0.054 | 1 | 1 |

**Supplementary Figures**

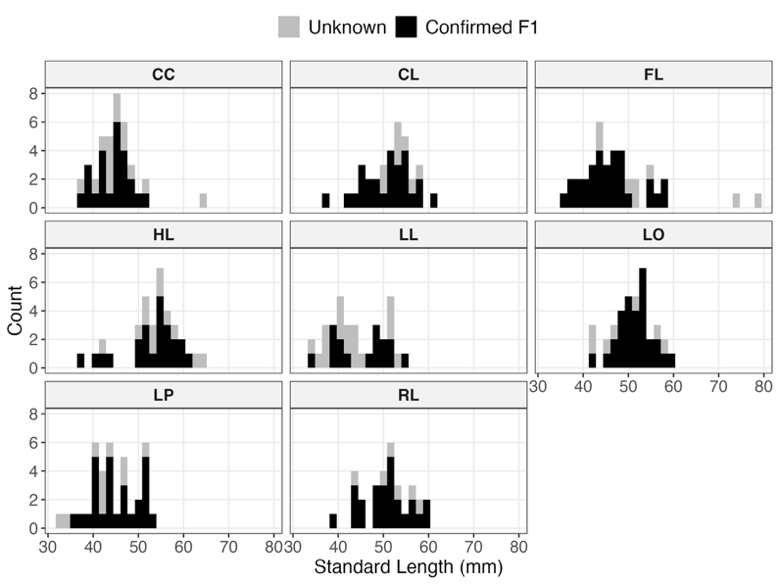

**Figure S1**: Age class inference for one-year post-introduction. Fish that were confirmed to be from the F1 generation (according to genotyping data) spanned the size distribution in all lakes, aside for three exceptionally large individuals (in CC and FL). These outliers were deemed to be from the parental generation and excluded from subsequent analysis while all remaining individuals were assumed to be from the F1 generation.

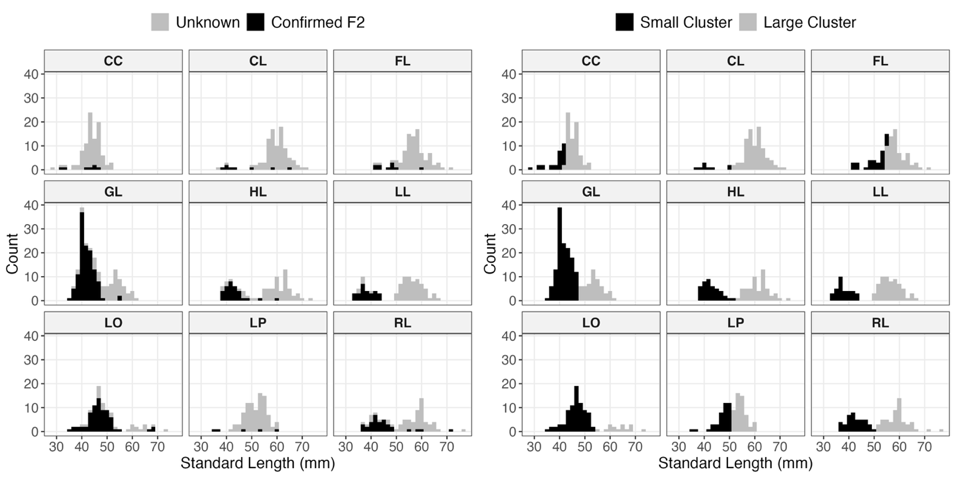

**Figure S2**: Age class inference for two years post-introduction. The left plot shows individuals that were confirmed to be from the F2 generation, according to genotyping data. The right plot shows results from k-means clustering by standard length within each lake. In CL, GL, HL, LL, LO, and RL, the size clusters appear to correspond to age classes, based on the distribution of the confirmed F2 individuals. To select fish from the F1 generation in these lakes, we retained only the individuals from the larger cluster and further removed any remaining confirmed F2 individuals. In CC, FL, and LP, there were very few confirmed F2 individuals and the size clusters were uninformative. In these cases, we simply omitted any confirmed F2 individuals and assumed the remaining individuals belonged to the F1 generation.

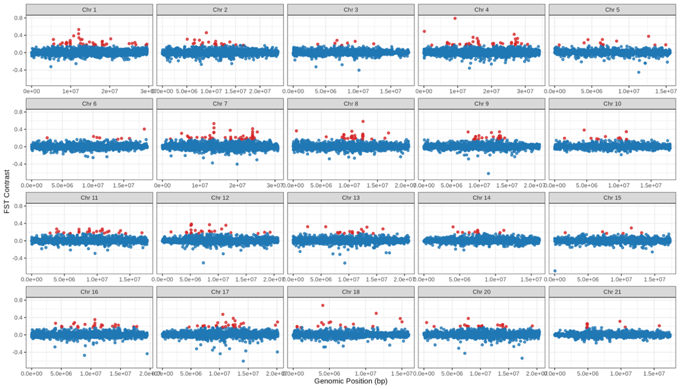

**Figure S3**: Ecotype F_ST_ contrast. The mean F_ST_ of between-ecotype comparisons minus the mean F_ST_ of within-ecotype comparisons in 10kb windows across all the autosomes. Red points are the putatively adaptive windows (top 1% outliers) that were used to subsequently calculate adaptive F_ST_ among populations.

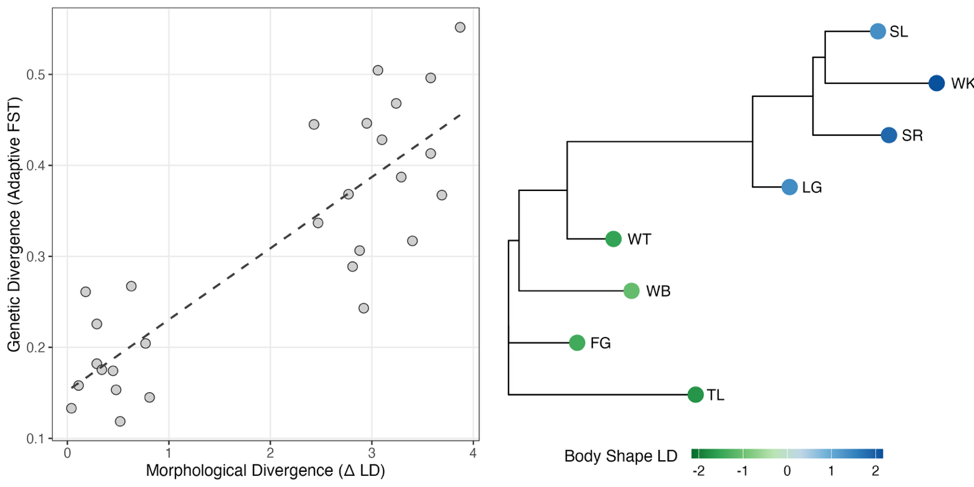

**Figure S4**: Validating adaptive F_ST_. Adaptive F_ST_ compared to morphological divergence among populations (left panel) and a neighbour joining tree of adaptive F_ST_ values (right panel), showing that the populations are clustered into ecotypes.

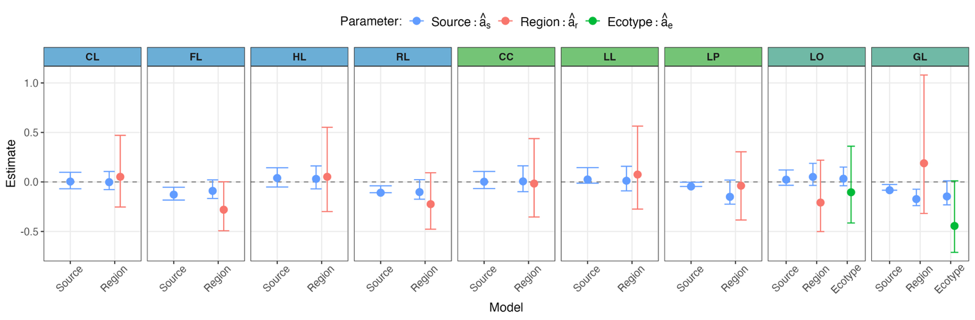

**Figure S5**: Assortment parameter estimates. Estimates of each parameter in each model with 95% confidence intervals. Source population models only include assortment by source population, while region and ecotype models include assortment by region and ecotype, respectively, as well as assortment by source population.

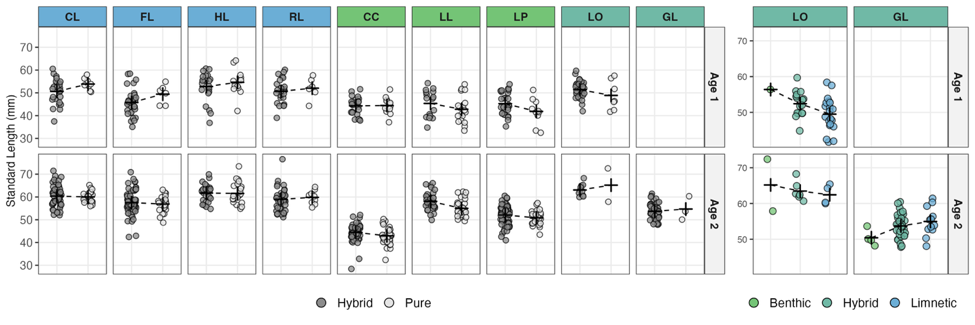

**Figure S6**: Selection on hybrid genotypes: Panels on the left compare standard length between pure and hybrid genotypes according to source population, within each lake and each age class, with dashed lines connecting the means. Panels on the left compare standard length between ecotypic genotypes (pure benthic, pure limnetic, and hybrids). Note that our entire sample of fish from GL came from two years post introduction, hence why the panels are empty for age 1.

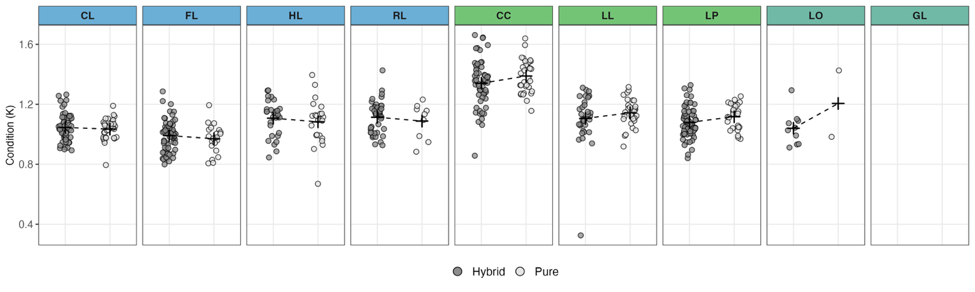

**Figure S7**: Selection on hybrid genotypes. Comparisons of condition index (K, length/mass^3^) between hybrid and pure genotypes in each lake, none of which were significant (Table S12). Note that mass data was only available for individuals collected two years post-introduction, thus all of these fish are age two.

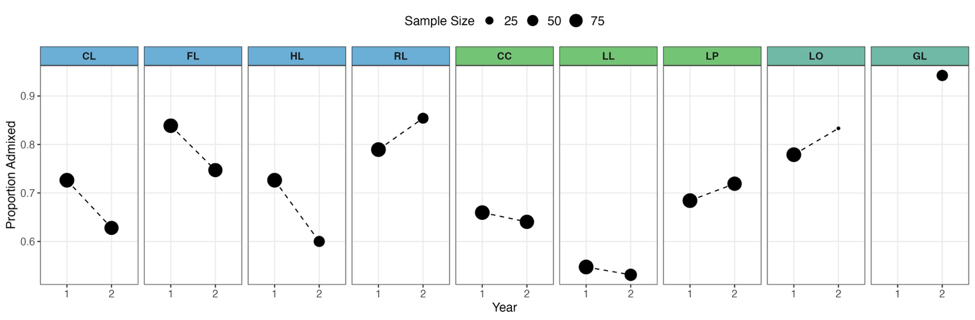

**Figure S8**: Proportion of hybrid individuals in each lake one year and two years after introduction. None of these comparisons were significant (Table S13).
